# ERK5 Signaling is Required for Type III IFN-mediated Mucosal Antiviral Responses

**DOI:** 10.1101/2024.05.21.595218

**Authors:** Hannah Bone, Dalia S. Natour, Matthew I. McFadden, Andrew Karp, Anandita Basu, Andrea Keller, Parker Denz, Patrick L. Collins, Maria M. Mihaylova, Jacob S. Yount, Adriana Forero

## Abstract

Type III interferons (IFNλ) are innate immune cytokines that limit viral replication and coordinate tissue repair through the induction of interferon stimulated genes (ISGs). This response must be tightly regulated to avoid excessive responses that result in the disruption of tissue barrier integrity or inefficient responses that allow for pathogen escape. Here we examine the contribution of Mitogen Activated Protein Kinase (MAPK) signaling on IFNλ-mediated antiviral activity. We find that extracellular-signal-regulated kinase 5 (ERK5), a poorly characterized member of the conventional MAPK family, potentiates the antiviral efficacy of IFNλ. Chemical inhibition and genetic targeting of ERK5 during IFNλ treatment of cells results in a decrease in ISG induction and impaired control of viral infections. This decrease in IFNλ antiviral efficacy in the absence of ERK5 kinase activity corresponded to lowered STAT1 phosphorylation, revealing a noncanonical role for ERK5 in STAT1 activation downstream of IFNλ. In contrast, type I IFN antiviral signaling is largely resistant to ERK5 modulation. Altogether, we identify ERK5 as a potentiator of STAT1 activation, ISG expression, and antiviral activity following type III IFN stimulation.

**SIGNIFICANCE:** Regulation of type III interferons (IFNλ) at mucosal barriers in response infection to mitigate viral replication and support barrier integrity. The specific mechanistic requirements for MAPK signaling to sustain IFNλ-mediated gene expression have remained elusive. Amongt the least characterized members of the MAPK family, the role of ERK5 in regulating host inflammatory responses has been hampered by off-target effects of kinase inhibitors. Here, we combine pharmacological and genetic approaches to specifically demonstrate that ERK5 promotes antiviral immunity in epithelial cells. Mechanistically, ERK5 enhances the activation of STAT1 in response to IFN stimulation to augment the transcription of IFN-stimulated genes. Our work demonstrates that therapeutic modulation of MAPK and IFN signaling pathway co-integration could distinguish between the protective and deleterious outcomes of IFN expression.

*One-sentence summary:* ERK5 potentiates IFN lambda responses.

## INTRODUCTION

Interferons (IFN) are cytokines induced upon pathogen infection or the loss of organelle integrity. Two IFN families, type I (IFNα/β) and type III (IFNλ) IFNs, signal through cognate IFN receptors (IFNα/β - IFNAR1 and IFNAR2; IFNλ – IFNLR1 and IL10R2) to activate canonical Janus kinase (JAK)-signal transducer and activator of transcription (STAT) signaling pathways necessary to drive IFN-stimulated gene (ISG) expression. Many of the hundreds of ISGs encode proteins with direct antiviral or antiproliferative activities that can inhibit pathogen replication and cell proliferation. Balancing of this innate immune response is critical to curb viral replication while supporting tissue repair responses and preventing uncontrolled cellular proliferation.

Despite having shared antiviral roles, it has been well established that IFNα/β and IFNλ have unique functions (1). In vivo studies support distinct roles for IFNα/β and IFNλ in the induction of inflammatory responses during infections (2–4). The mechanistic insights into the shared and distinct functions of IFNα/β and IFNλ have been derived primarily from delineating differences in the activation of canonical JAK/STAT signaling and IFN regulatory factors (IRF) which distinguish their ability to induce the expression of antiviral and inflammatory ISGs (5, 6). On the other hand, it is known that IFNs can elicit distinct responses across cell types. Recent work demonstrated that IFNλ can promote intestinal barrier integrity by eliciting distinct gene expression programs in intestinal stem cells, found in crypts, relative to terminally differentiated epithelial cells (7). Although the mechanistic underpinning of these differential responses is not fully understood, it is possible that highly regulated expression of IFNλ receptors could account for the robustness of the transcriptional programs induced across different cells types in the epithelium (1, 6, 8). Whether additional molecular cues tune signal transduction and gene transactivation in response to IFN stimulation remains incompletely studied.

Mitogen-activated protein kinases (MAPK) pathway activation is triggered in response to cytokines, growth factors, stress signals, and pattern recognition receptor engagement associated with the activation of ISGs (9–11). Our previous work revealed that that cells stimulated with IFNλ elicit gene expression profiles that are associated with the activation of MAPK signaling (6). This enrichment was not observed when cells were stimulated with IFNβ. Other studies have reported a differential MAPK dependency in setting the threshold of antiviral potency between IFNα/β and IFNλ (12). They demonstrated that the induction of an antiviral state in response to IFNλ could be dampened by chemical inhibition of the conventional MAPK pathways involving p38, JNK, and ERK1/2 (12) and that the cellular antiviral state elicited by IFNα/β stimulation was less sensitive to the loss of MAPK activity. These studies support the relevance of signal co-integration between the MAPK and IFN pathways and warrant futher investigation to determine how this tunes the kinetics and magnitude of the IFN responses to dictate IFN functions.

A lesser studied member of the conventional MAPK family, Extracellular-signal-Regulated Kinase 5 (ERK5 or BMK-1) (13) is similarly activated by stress and growth factor stimulation. ERK5 activation regulates cell cycle progression and has been shown to promote proliferation of malignant cells (14). ERK5 is also essential for angiogenesis (15) and tumor clearance (16, 17). As such, pharmacological perturbation of ERK5 has become an attractive approach for cancer therapy. However, the extent to which inhibition of ERK5 could impair host protective responses, such as resistance to viral infection, is less clear. The activation of Unc-51-like kinase 1 (ULK1) by type II IFN (IFN_γ_) is required for antiviral protection (18). While this study suggested that ULK1 induces ERK5 to act as a potential regulator of IFN_γ_ responses, the mechanisms by which ULK1/ERK5 regulates gene expression in this context remain elusive. Importantly, whether such dependency is conserved across IFN families is unclear.

Here we explore the requirement for ERK5 (encoded by *Mapk7*) activity in sustaining ISG expression downstream of IFNλ receptor activation in murine intestinal epithelial cells. We demonstrate that inhibition of ERK5 kinase activity or *Mapk7* gene ablation impairs ISG transcription and protein expression. This decrease in effector gene expression rendered IFNλ-treated cells susceptible to viral infection. Mechanistically, ERK5 is required to promote the activation of signal transducer and activator of transcription 1 (STAT1). Interestingly, inhibition of ERK5 did not drastically impair ISG induction or antiviral protection by IFNβ stimulation. This study demonstrates the preferential reliance of IFNλ on integration of MAPK signaling to promote robust ISG expression.

## RESULTS

### ERK5 kinase activity enhances type III IFN mediated ISG expression

Although ERK5 is thought to be ubiquitously expressed, whether the distribution and activation of ERK5 varies across tissues is poorly understood. We examined total ERK5 protein expression and activation (T218/Y220 phosphorylation) in the small intestine (SI), an organ composed of heterogenous epithelial cell types and distinct regional function. We found ERK5 to be constitutively active across the small intestine with total ERK5 expression highest in crypts relative to villi (Figure 1A). We also noted that proximal SI ERK5 expression was higher than that observed at distal sites. As a control, we measured GATA4 protein expression which as a regional defining marker (19, 20). Expression of GATA4 was expectedly higher in the proximal SI relative to the distal SI (Figure 1A). Previous reports have demonstrated differences in the qualitative and quantitative response to IFNλ stimulation across cell types enriched in crypts and those derived from villi (7). To determine whether ERK5 kinase activity could influence interferon-stimulated gene (ISG) expression in response to IFNλ, we pre-treated immortalized murine SI epithelial cells (muINTEPI) with a selective ERK5 kinase inhibitor, XMD8-85, for two hours prior to stimulation with recombinant murine IFNλ. Total RNA was harvested 24 hrs post IFNλ stimulation and genome-wide mRNA expression analysis was conducted. We observed that ERK5 kinase inhibition reduced the number of differentially expressed genes (DEG; LFC |0.26|, p-value adj. 0.05) relative to vehicle control treated cells (Figure 1B, Figure S1A). Analysis of the magnitude of DEG expression revealed that ERK5 kinase activity led to a significant global decrease across ISGs (Figure 1C, Supplementary Table 2). Gene set enrichment of biological processes inferred a decrease in the antiviral effector genes that are induced by IFNλ stimulation (Figure S1B). We examined the protein expression of the identified DEF by Capillary-LC/MS/MS. We observed that ERK5 inhibition generally inhibited protein accumulation relative to DMSO-treated cells (Figure 1D, Supplementary Table 3). These proteins included the transcription factors STAT1, IRF9, and IRF7 (Figure S1D), which are positive regulators of ISG expression following IFN stimulation (1).

**Figure 1.**
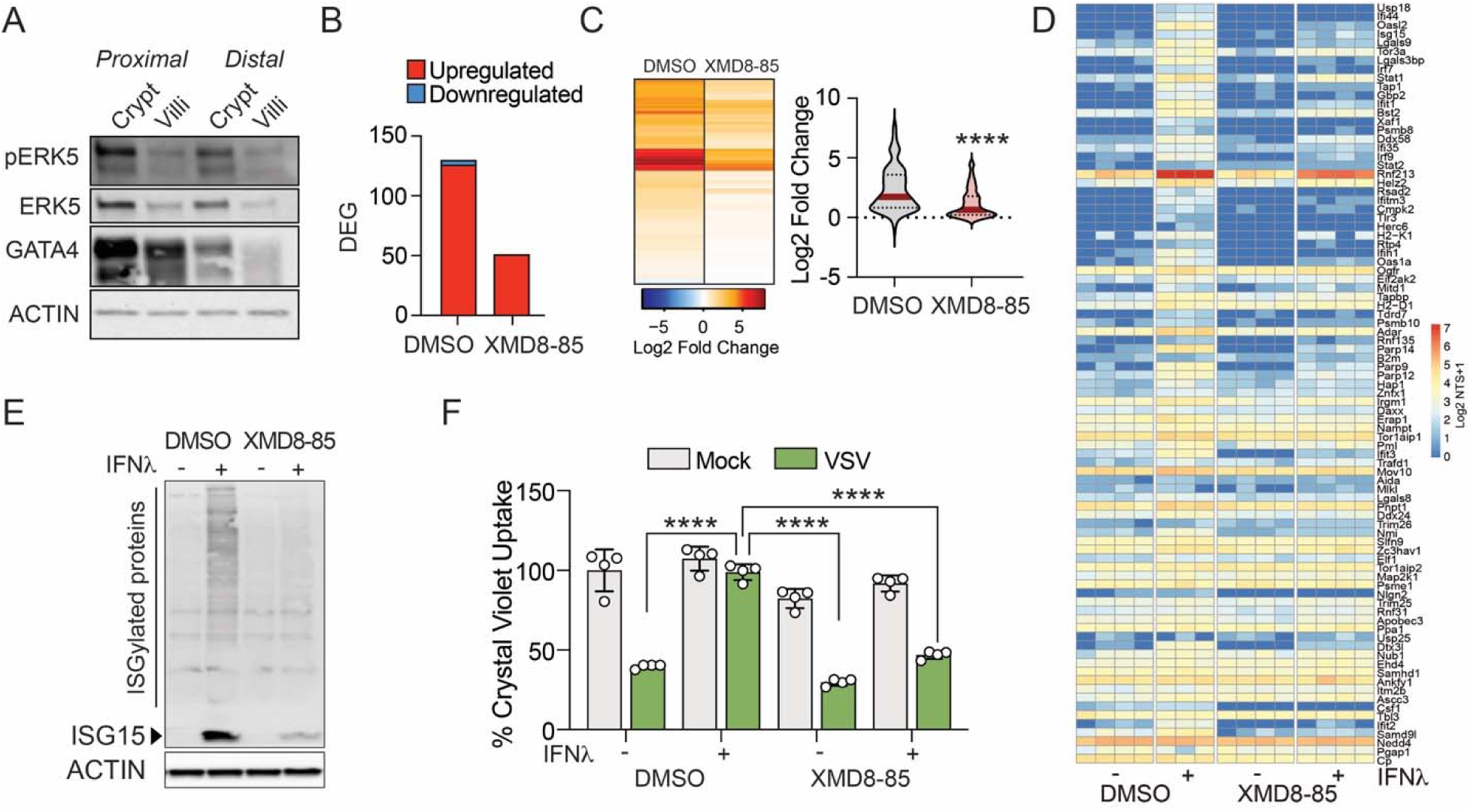
ERK5 kinase activity is required to support type III IFN-mediated ISG expression. Expression of activated ERK5 kinase in murine small intestinal crypts and villi. (A) Western blot expression analysis of phosphorylated ERK5 (T218/Y220), total ERK5, GATA4, and Actin across crypts and villi obtained from proximal and distal small intestinal sites. (B) Chemical inhibition of ERK5 kinase activity (XMD8-85; 10 μM) inhibits type III IFN (IFNλ)-mediated induced transcriptional changes in muINTEPI cells. Bar graph represents the number of differentially expressed genes (DEG) (LFC |0.26|; adj p-value 0.05) following IFNλ stimulation in cell pre-treated with either vehicle (DMSO) or XMD8-85. Color represents genes upregulated (red) or inhibited (blue) by IFNλ relative to DMSO or XMD8-85, mock-treated cells. (C) Relative expression heatmap of the union of 131 DEG (left; Supplementary Table 2). Violin plot represents the expression 131 ISGs following IFNλ treatment (50 ng/ml) for 24 h (right). Median gene expression is denoted by red line and black dotted lines indicate confidence intervals. Statistical significance was derived using student’s t test. (D) Protein expression of 83 DEG detected by LC-MS/MS in cells pretreated with DMSO or XMD8-85 followed by IFNλ stimulation across individual samples. For each condition there are 3-4 individual replicates. Color indicates the log2 transformed normalized total spectra + 1. (E) Western blot analysis of ISG15 protein expression. Whole cell lysates were prepared from cells treated with XMD8-85 (10 μM) for 2 h prior to stimulation with IFNλ treatment (50 ng/ml) for 24 h. Data representative of three individual experiments. (F) muINTEPI monolayers were pre-treated with (XMD8-85; 10 μM) 2 hrs prior to IFNλ stimulation (100 ng/ml) for 24 hrs. Media was replaced with fresh growth medium without inhibitor or cytokine prior to infection with VSV-GFP (MOI = 3) for 18 hrs. Cells were fixed and stained for cellular crystal violet uptake. Relative uptake calculated to DMSO, uninfected cell absorbances (relative value 100%).

We confirmed our transcriptional and proteomic observations by targeted assessment of gene expression. ERK5 kinase inhibition resulted in a decrease in IFNλ-mediated *Isg15* and *Ifit1* mRNA induction in independent experimental sample sets (Figure S1C). Similarly, we confirmed our proteomics data by examining the effect of ERK5 kinase inhibition on ISG15 protein expression by western blot. Following IFNλ3 stimulation, ISG15 protein expression was inhibited by ERK5 kinase inhibition compared to DMSO treatment (Figure 1E). Consequently, XMD8-85 pre-treatment led to a decrease in the protective efficacy of IFNλ against the cytolytic activity of Vesicular Stomatitis Virus (VSV-GFP) in infected cells (Figure 1F). Together, these results suggest that ERK5 kinase activity is necessary for robust transactivation of ISGs and effector protein expression to mediate the antiviral efficacy of type III IFNs.

### Evaluation of immunomodulatory efficacy of ERK5 small molecule inhibitors

We next examined the dose dependent effect on ISG repression elicited by ERK5 kinase inhibition. Pre-treatment with XMD8-85, with as low as 1 μM, led to a significant inhibition of *Ifit1* (left) and *Isg15* (right) mRNA expression at 24 hrs post IFNλ stimulation (Figure 2A) and suppression of ISG mRNA levels was observed in a dose dependant manner. Although small molecule inhibitors of ERK5 show selectivity over other kinases, they are known to have off-target effects that could confound our data interpretation (21). We tested the effect of ERK5 inhibition with JWG-071 and noted a significant dose-dependent decrease in *Ifit1* (left) and *Isg15* (right) mRNA expression in inhibitor pre-treated cells relative to DMSO treated cells (Figure 2B). Lastly, we evaluated the effect of AX15836, a potent and highly selective ERK5 inhibitor that lacks bromodomain inhibitory activity, on the induction of antiviral gene expression following IFN stimulation (Figure 2C). As observed with XMD8-85 and JWG-071, treatment with 0.5 μM of AX15836 resulted in a decrease in ISG mRNA levels in response to IFNλ stimulation. This inhibitory effect was more prominent on *Isg15* than *Ifit1* mRNA expression and *Isg15* expression was significantly decreased in ERK5 inhibitor treated cells, relative to vehicle control cells. Together, these data suggest that small molecule inhibitors of ERK5 kinase decrease the transactivation of ISGs after IFNλ stimulation.

**Figure 2.**
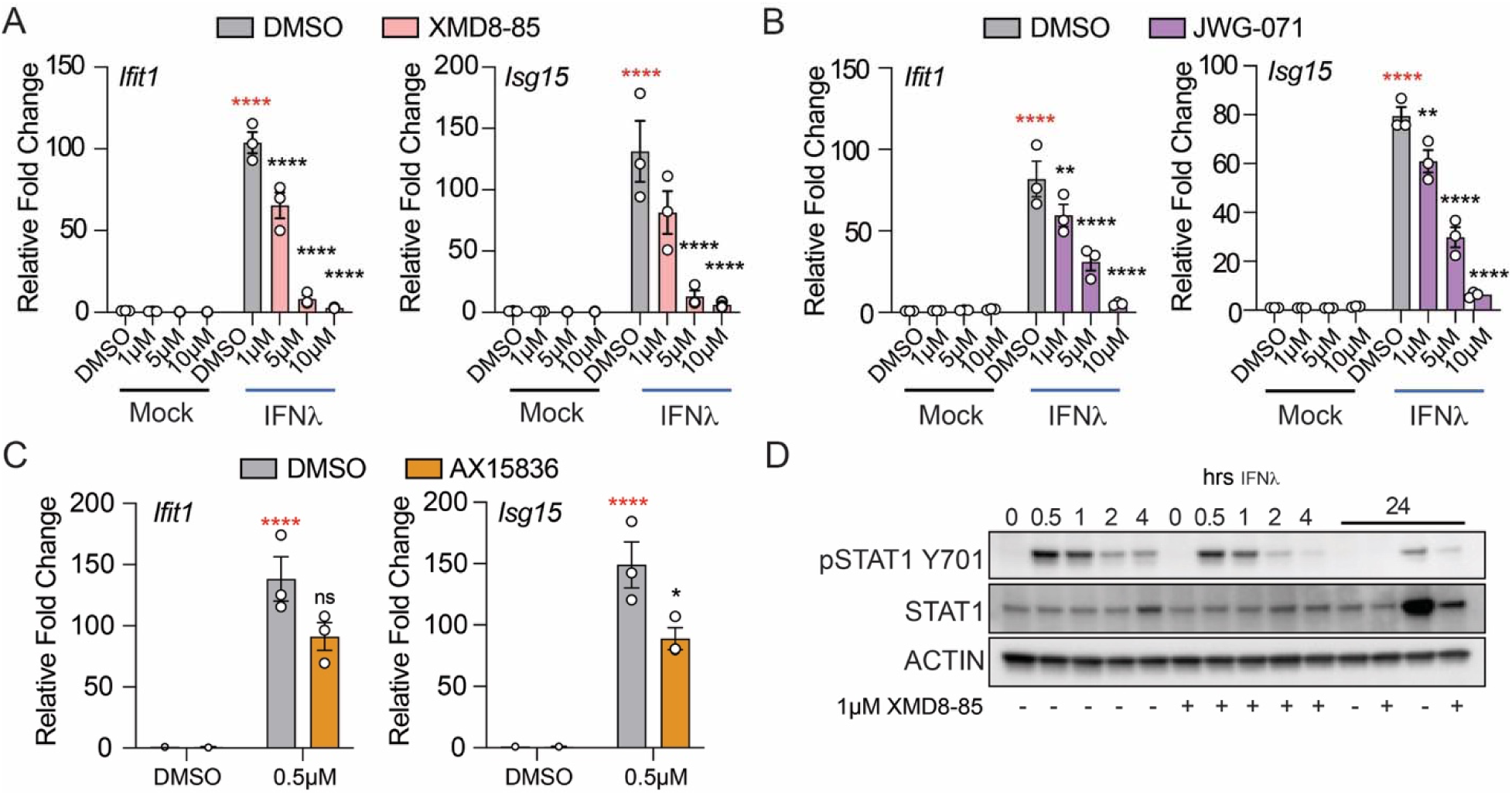
Dose-dependent ISG decreases in responses to ERK5 kinase inhibition. (A) Dose response to XMD8-85. Murine SI cells (muINTEPI) cells were treated with increasing doses of XMD8-85 or vehicle control (DMSO) for 2 hrs prior to stimulation with murine recombinant IFNλ (50 ng/ml) for 24 hrs. Relative expression of *Ifit1* (left) and *Isg15* (right) mRNA was normalized to the expression of the house keeping gene *Chmp2a* and DMSO treated, mock-stimulated cells (relative value 1). Data is representative of the average of three individual replicates and error bars represent ±SEM. Asterisk represent significance relative to mock treated cells (red) or DMSO-treated, IFN stimulated cells (black) as determined by two-way ANOVA. (B) Dose response to ERK5 kinase inhibitor, JWG-071. Cells were pre-treated with inhibitor and/or IFNλ as indicated above. Relative expression of *Ifit1* (left) and *Isg15* (right) mRNA was normalized to the expression of the house keeping gene *Chmp2a* and DMSO treated, mock-stimulated cells (relative value 1). Data is representative of the average of three individual replicates and error bars represent ±SEM. Asterisk represent significance relative to mock treated cells (red) and DMSO-treated, IFNλ stimulated cells (black) as determined by two-way ANOVA. (C) ISG modulation by ERK5 inhibitor, AX15836. Cells were pre-treated with inhibitor, vehicle control, and/or IFNλ as indicated above. Relative expression of *Ifit1* (left) and *Isg15* (right) mRNA was normalized to the expression of the house keeping gene *Chmp2a* and DMSO treated, mock-stimulated cells (relative value 1). Data is representative of the average of three individual replicates and error bars represent ±SEM. Asterisk represent significance relative to mock treated cells (red) and DMSO-treated, IFNλ stimulated cells (black) as determined by two-way ANOVA. (D) Activation of STAT1 following IFNλ treatment. Whole cell lysates from muINTEPI cells pre-treated with XMD8-85 (1 μM) followed by IFNλ stimulation for the indicated time points were probed for expression of phosphorylated STAT1 (Y701), total STAT1, and Actin.

To address the molecular mechanism of ISG regulation, we measured the effect of ERK5 inhibition on the activation of STAT1 following IFN stimulation. Inhibition of ERK5 kinase activity with XMD8-85 suppressed the activation of STAT1, as determined by phosphorylation at tyrosine 701 (Y701). The decrease in STAT1 Y701 phosphorylation was most prominent at 1 to 4 hrs post IFNλ stimulation. At 24 hrs post stimulation, both total STAT1 and STAT1 Y701 protein expression was blunted in XMD8-85 pre-treated cells relative to vehicle-treated, IFN stimulated cells (Figure 2D). These data indicate that ERK5 kinase activity is necessary to promote the activation of STAT1 downstream of the IFNλ receptor to promote robust transactivation of ISGs.

### Genetic perturbation of ERK5 confirms its antiviral regulatory role

A major challenge in the functional characterization of ERK5 is that phenotypes observed using kinase inhibitors are poorly recapitulated by genetic ablation (22). To better define the specific role of ERK5 in the regulation of IFN responses, we generated *Mapk7*- knockout (KO) muINTEPI cells by CRISPR-Cas9 genome editing. Clones were evaluated to assess ERK5 depletion and expression of conventional MAPK family members, p38, JNK, and ERK1/2 (Figure S2A). Additionally, clones were selected for downstream analysis based on equivalent expression of the high-affinity type III IFN receptor subunit, *Ifnlr1,* and the type I IFN receptor, *Ifnar1* (Figure S2B). Wild-type (WT) and *Mapk7* KO clones with comparable expression of other MAPK (Figure 3A) were further screened to examine the requirement for ERK5 expression and function in the induction of ISGs following IFNλ stimulation.

**Figure 3.**
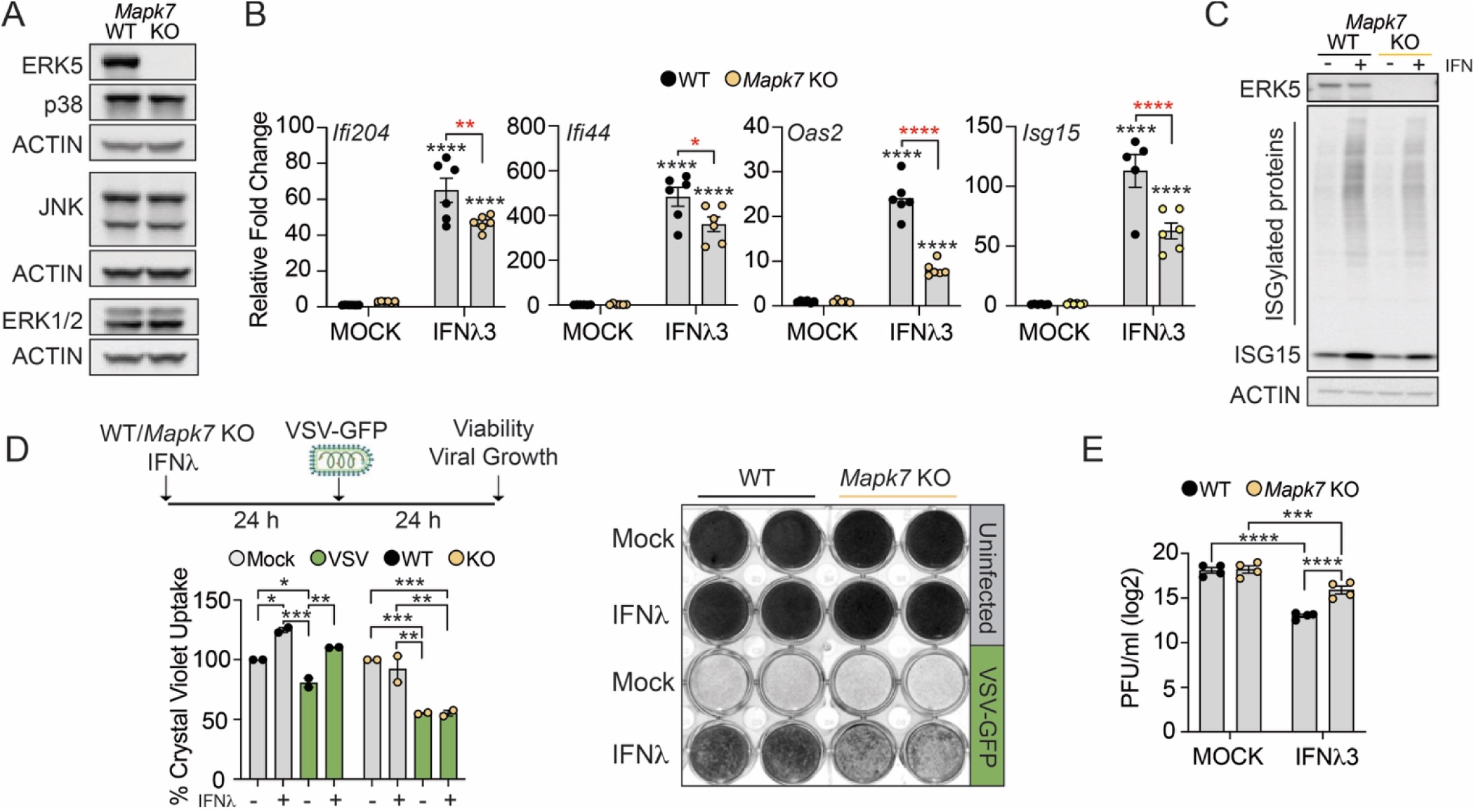
Loss of ERK5 phenocopies the inhibition of kinase activity. (A) Validation of ERK5/*Mapk7* targeting in muINTEPI cells. Cells were transduced with Cas9 and non-targeting sgRNA (WT) or *Mapk7*-targeting sgRNA. Western blot analysis of total ERK5, ERK1/2, p38, JNK, and Actin protein expression. (B) Requirement of ERK5 expression for ISG induction. Wild type or *Mapk7* null muINTEPI cells were treated with IFNλ (50 ng/ml) stimulation for 24 h prior to total RNA extraction. Bar graphs represent the average expression of *Ifi204*, *Ifi44*, *Oas2,* and *Isg15* mRNA across six independent experiments and error bars represent ±SEM. Asterisks represent p-value of significance as determined by two-way ANOVA. (C) Western blot analysis of ISG15 protein expression following IFN treatment of WT and muINTEPI cells. Cells were treated IFNλ (100 ng/ml) for 24 hrs prior to preparation of whole cell lysates. Data representative of 3 independent experiments. (D) Cellular crystal violet uptake following ERK5 kinase modulation, IFN treatment and viral challenge as previously described. Bar graphs represent normalized (100%) average uptake across 2 independent experiments relative to vehicle, mock-treated samples. Asterisks represent p-value of significance as determined by two-way ANOVA (left). Representative image of cell culture plates (right). (E) Quantification of infectious VSV production in WT and *Mapk7* KO cells as determined by plaque assay. Bar graphs represent average log2 transformed viral titers across 4 independent experiments and error bars represent ±SEM. Asterisks represent p-value of significance as determined by two-way ANOVA.

Relative to WT cells, the deletion of ERK5 resulted in a significant decrease in the magnitude of ISG induction (*Ifi204, Ifi44, Oas2, Isg15*) at 24 hrs post IFN stimulation relative to WT cells (Figure 3B). As observed in ERK5 inhibitor treated cells, the decrease in ISG mRNA was accompanied by a decrease in unconjugated ISG15 protein levels and reduced protein ISGylation in cells lacking ERK5 expression (Figure 3C). We examined the functional consequences of *Mapk7* deletion on IFNλ antiviral immunity using a crystal violet uptake assay (Figure 3D). Virus infection of mock-stimulated cells led to a significant decrease in crystal violet uptake in both genotypes (Figure 3D). Treatment with IFNλ protected cells from VSV-mediated cytotoxicity. However, the loss of *Mapk7* abrogated IFNλ protective responses relative to the that observed in WT cells. *Mapk7* deficiency did not alter dye uptake in either uninfected mock or IFNλ-stimulated cells. To corroborate these results, we quantified infectious virus production in in VSV-infected WT and *Mapk7* KO cells in the presence or absence of IFNλ stimulation. Consistent with the crystal violet assays, comparable viral growth was observed in mock-stimulated, VSV-infected cells. Although IFN pre-treatment led a significant decrease in viral replication both WT and KO cells, VSV replication was significantly higher in *Mapk7* KO cells relative to WT cells following IFN treatment (Figure 3E, Figure S2C). Together, our genetic data support that ERK5 expression is necessary to promote efficient IFNλ-mediated antiviral immunity.

### Conserved inflammatory requirement for ERK5 kinase activity across epithelial cells

We examined a potential conservation for ERK5 antiviral dependency across epithelial cell lines. We pre-treated A549 human lung epithelial cells with increasing doses (1μM – 10μM) of XMD8-85 prior to human recombinant IFNλ3 stimulation. As observed in murine SI cells, ERK5 kinase inhibition resulted in a dose-dependent decrease in antiviral effector protein expression (i.e. RIG-I, IFIT3, IFIT3, and ISG15) (Figure 4A). IFNλ stimulation of A549 cells significantly decreased VSV infectivity relative to the infection levels observed in DMSO-treated, unstimulated cells (Figure 4B). Treatment with either 1 μM or 10 μM of XMD8-85 restored VSV infectivity to levels comparable to those observed in DMSO-treated, unstimulated cells (Figure 4B). We also measured IFN-mediated antiviral protection against a respiratory pathogen, influenza A virus (IAV). IFNλ stimulation reduced IAV infectivity of A549 cells (Figure 4C). We noted a dose-dependent increase in the number of IAV-infected cells (Figure 4C) in IFNλ and XMD8-85 co-stimulated cells compared to IFNλ and DMSO-treated cells. ERK5 inhibition alone did not impact the infectivity of either VSV or IAV. These data demonstrate that ERK5 kinase activity is predominantly necessary to establish a robust pan-antiviral state following IFN sensing.

**Figure 4.**
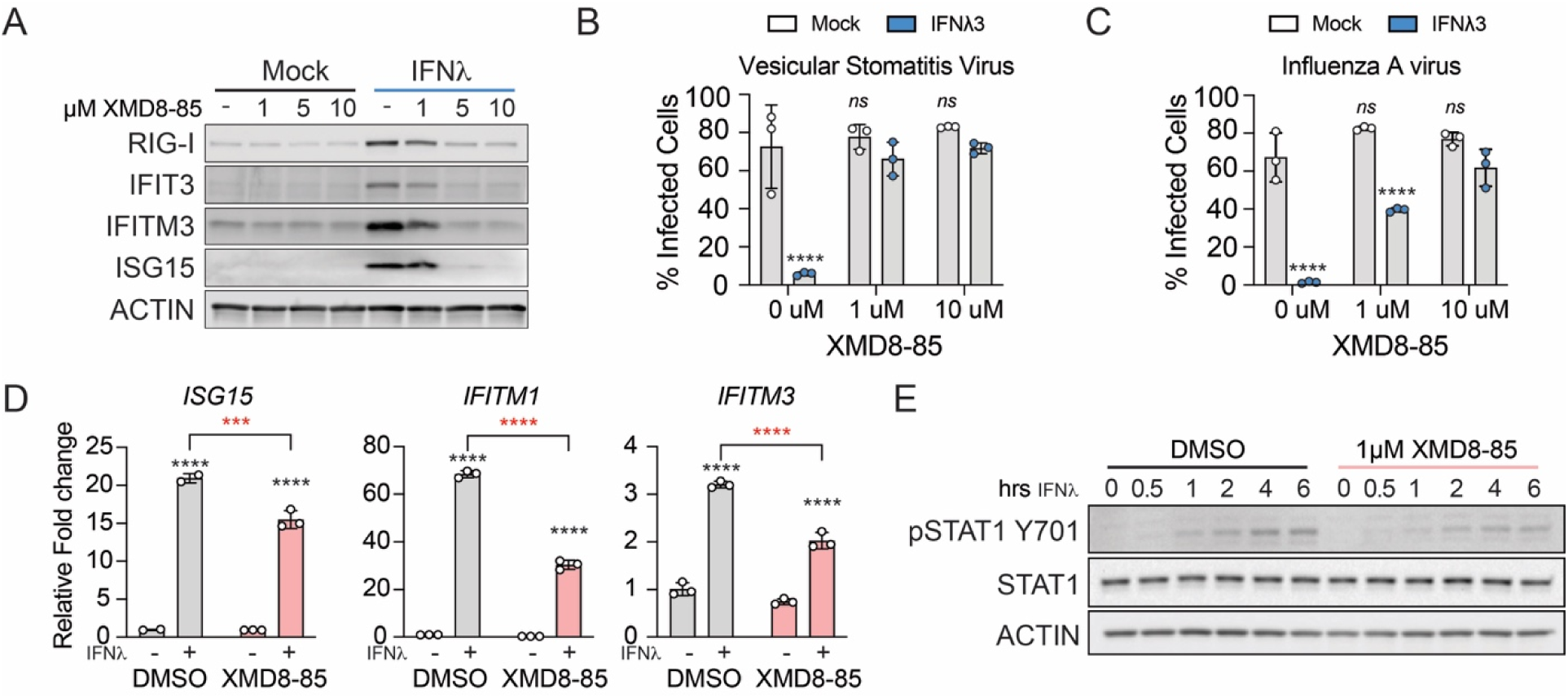
The requirement for ERK5 kinase activity is conserved across cell types. (A) Induction of ISG protein expression in human airway cell line, A549, treated with human recombinant IFNλ3 (50 ng/ml). Total expression of RIG-I, IFIT3, IFITM3, ISG15, and Actin was measured using western blot. (B) Dose dependent effect of ERK5 kinase inhibition on viral replication. A549 cells were pretreated with 1 μM or 10 μM XMD8-85 before co-stimulation with IFNλ prior to infection with vesicular stomatitis virus (VSV-GFP; MOI 0.01). Percent infected cells was determined by flow cytometric analysis of GFP positive cells. Bar graph represents the average cell infection across three independent experiments. Statistical significance was assessed using two-way ANOVA. (C) A549 cells were pretreated with 1 μM or 10 μM XMD8-85 before co-stimulation with IFNλ prior to infection with Influenza A Virus (PR8-GFP; MOI 5). Percent infected cells was determined by flow cytometric analysis of GFP positive cells. Bar graph represents the average cell infection across three independent experiments. Statistical significance was assessed using two-way ANOVA. (D) Effect of ERK5 kinase inhibition on immortalized human hepatocyte cells. Bar graphs represent *ISG15*, *IFITM1,* or *IFITM3* mRNA expression after pre-treatment of PH5CH8 cells with DMSO or 1 μM XMD8-85 2 hrs prior to IFNλ (50ng/ml) stimulation. Error bars represent ±SD. Statistical significance was assessed using two-way ANOVA. Data is representative of 3 independent experiments. (E) Activation of STAT1 in the presence of ERK5 inhibitors. Whole cell lysates from PH5CH8 cells pre-treated with XMD8-85 (1 μM) or DMSO followed by IFNλ stimulation (50ng/ml) for the indicated time points were probed for expression of phosphorylated STAT1 (Y701), total STAT1, and Actin.

We then examined ERK5 kinase requirement in the ISG response to IFNλ in human immortalized hepatocytes, PH5CH8. Cells were pre-treated with XMD8-85 prior to IFNλ stimulation. Inhibition of kinase activity resulted in a significant decrease in the induction of ISG mRNA (i.e. *IFITM1, ISG15, STAT1*) relative to vehicle-control treated cells (Figure 4D). Consistent with our mechanistic observations in murine intestinal epithelial cells, the inhibition of ERK5 led to a decrease in STAT1 activation (Y701 phosphorylation) proximal to IFNλ stimulation (Figure 4E). The loss of ERK5 kinase activity did not impact the baseline expression of total STAT1. Overall, these data suggest that ERK5 kinase activity supports the antiviral efficacy of IFNλ across species and cell types likely through the conserved positive regulation of STAT1 phosphorylation.

### ERK5 kinase dependency predominant for type III IFN-mediated antiviral efficacy

Previous work found that type III IFNs have a preferential dependency on MAPK signal integration to promote robust antiviral protection (12). We incubated muINTEPI cells with 1 μM XMD8-85 2hrs prior to IFN stimulation for 24 hrs. As previously observed, the induction of expression of ISG15 protein in response to IFNλ stimulation was decreased in inhibitor-treated cells relative to vehicle control treated cells (Figure 5A). On the other hand, ISG15 protein induction by type I IFN was not affected by ERK5 kinase activity as expression levels were comparable between vehicle and inhibitor treated cells (Figure 5B). To evaluate the functional consequence of the observed differences in sensitivity to ERK5 modulation, we examined the impact of ERK5 kinase inhibition on the antiviral protective efficacy of IFNλ. As previously described, we pre-stimulated cells with XMD8-85 for 2 hrs prior to co-stimulation with IFNλ for 24 hrs. Both exogenous IFN and chemical inhibitors were removed prior to viral challenge for 24 hrs (Figure 5C). We leveraged the expression of green fluorescent protein within VSV-GFP infected cells (Figure 5D) to determine the impact of ERK5 inhibition on type I and III IFN-mediated antiviral protection at the single cell level. While XMD8-85 did not inhibit viral infectivity, treatment with either murine recombinant IFNβ or IFNλ led to a significant reduction in the proportion of virus positive cells (Figure 5E). ERK5 inhibition partially abrogated IFNλ mediated antiviral protection, but did not affect the reduction of infectivity induced by IFNβ pre-incubation. Of note, neither IFNλ stimulation nor XMD8-85 affected viability of non-infected cells (Figure 5F). The combination of XMD8-85 and type I IFN lead to a small, but significant increase in cell death (Figure 5F). Our data indicate that type I and III IFN signaling are differentially dependent on ERK5 kinase activity.

**Figure 5.**
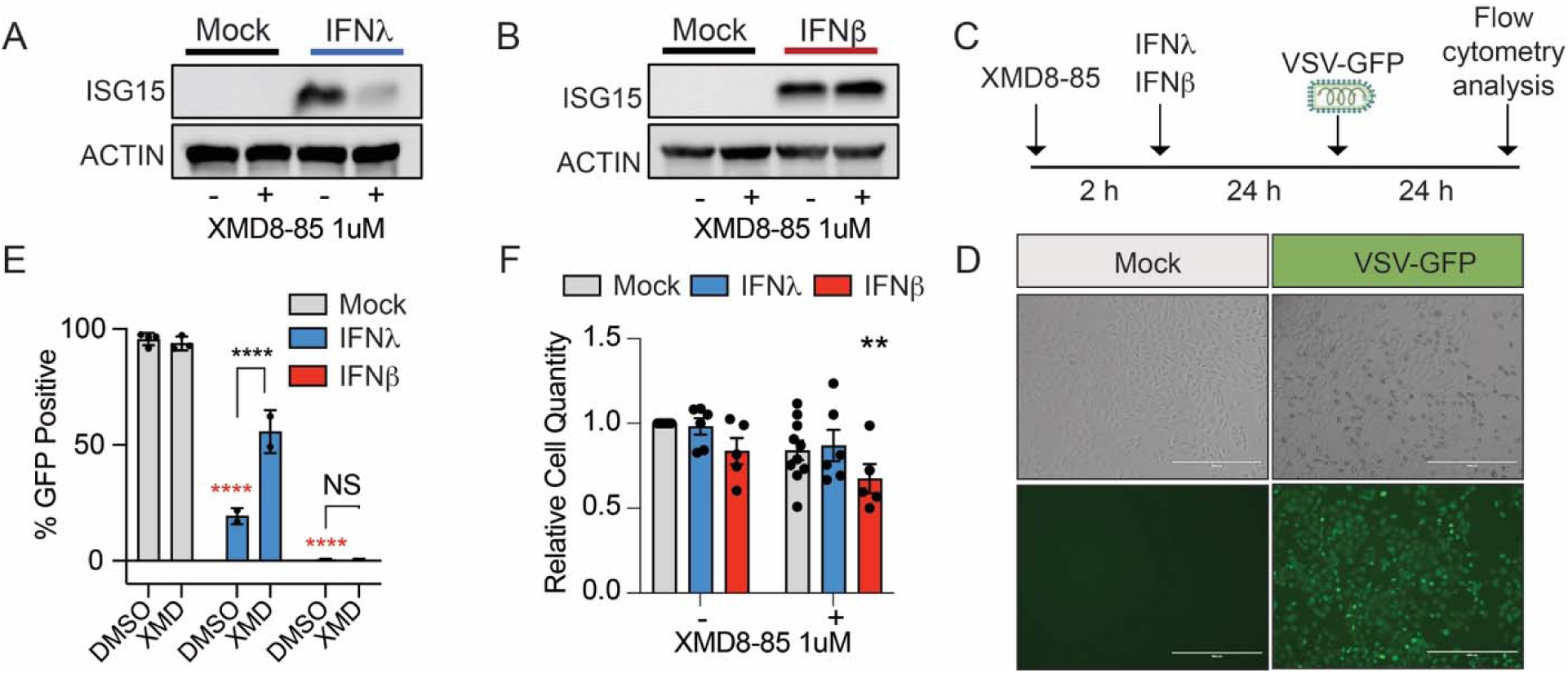
ERK5 kinase activity is predominantly required for type III IFN, but not type I IFN mediated antiviral protection. Western blot analysis of ISG15 protein expression in muINTEPI cells pre-treated with XMD8-85 prior to stimulation with (A) recombinant IFNλ3 (50 ng/ml) or (B) IFNβ (25 IU/ml). Data is representative of 3 independent experiments. (C) Schematic representation of experimental design to assess effect of ERK5 kinase activity on IFN induced antiviral responses. muINTEPI cells were pre-treated with XMD5-85 (1 μM) for 2 hrs prior to recombinant IFNβ (25IU/ml) or (IFNλ3 (50 ng/ml) stimulation for 24 h. Cells were then infected with VSV-GFP (MOI 0.01) for an additional 24 hrs prior to analysis of single cell green fluorescence by microscopy or flow cytometry. (D) Micrographs of VSV-GFP infected muINTEPI cells at an MOI 0.01. Green fluorescence signal was captured 24 h post infection by epifluorescent microscopy. (E) ERK5 inhibition diminishes antiviral protection elicited by type III IFN. Bar graphs represent the number of GFP-positive cells detected by flow cytometry following infection. Red asterisks indicate statistical significance relative to mock-treated DMSO-treated samples. Black asterisks indicate statistical significance between IFN treated groups as determined by two-way ANOVA. Data is average of 2-3 independent replicates. (F) Cellular viability following ERK5 kinase inhibition and IFN treatment. Bar graphs represent the relative cell quantity as determined by trypan blue exclusion assay and error bars represent ±SEM. Each data point indicates an independent experiment. Data was normalized to vehicle-treated, uninflected cells (Value 1).

## DISCUSSION

In this study, we have identified ERK5 as a novel regulator of type III IFN mediated antiviral protection. Pharmacological inhibition of ERK5 kinase activity resulted in a decrease of ISG transcription and a concomitant decrease in protein expression. Mechanistically, we found that inhibition of ERK5 kinase resulted in muted phosphorylation of STAT1, which is required for the formation of active transcription regulatory complexes that positively regulate ISG transcription (23). Of note, these phenotypes were recapitulated across mouse intenstinal cells and human airway epithelial and hepatocyte cell lines indicating a conserved requirement for ERK5 in the induction of ISGs across cell types and species.

The development of ERK5 kinase with selectivity for ERK5 over other non-target kinases has been a challenge for the field. To account for potential off-target effects, we corroborated findings derived from XMD8-85 inhibition using additional ERK5 kinase inhibitors. Treatment with either JWG-071 or AX15836 largely recapitulated the trend of decreased ISG induction that was observed in XMD8-85 treated cells. We further validated the role of ERK5 in IFN signaling by genetically ablating *Mapk7* and measuring the functional outcomes of IFN stimulation. We noted concordance in the effects of chemical inhibition of ERK5 versus ERK5 deletion in terms of decreased ISG expression and STAT1 phosphorylation. These data confirmed the specific requirement of ERK5 for the induction of antiviral programs.

Our data indicates that the establishment of an antiviral state in response to IFNλ was more reliant on ERK5 kinase activity than the antiviral protection conferred by IFNβ stimulation. Indeed, ISG protein expression induced by type I IFN was not significantly impaired by inhibition of ERK5 kinase activity. One possible explanation for this difference could be the magnitude of STAT1 induction downstream of IFN receptors. IFNλ affinity maturation studies have demonstrated that increasing receptor affinity enhances the kinetics of STAT1 phosphorylation and the antiviral potency of type III IFNs (24). In line with these observations, we have shown that STAT1 phosphorylation is limited by low IFNλ receptor abundance, which can be overcome IFNLR1 overexpression (6). On the other hand, STAT1 activation by type I IFNs is far more robust and thus could be resistant to changes in ERK5 activity (6, 25) could explain the decreased reliance on MAPK signal integration for the establishment of robust antiviral responses.

Signal transduction through the IFNα/β and IFNλ receptors is mediated by JAKs, non-receptor tyrosine kinases that facilitate receptor docking and phosphorylation of STAT1 and STAT2 (1). Phosphorylated STAT1 and STAT2 dimerize and bind IRF9 to activate ISG expression by binding to ISRE in the promoter of target genes. While JAK1 is essential for gene induction in response IFNα/β and IFNλ (26–29), the TYK2 IFN-mediated STAT activation appears to be more cell-dependent. Patients and mice lacking TYK2 expression have compromised JAK/STAT phosphorylation in response to IFNα/β stimulation (30–32). Although the response to IFNλ stimulation is partially impaired in TYK2-null patient-derived cells, intranasal inoculation with IFNλ protects *Tyk2*^−/−^ mice from IAV disease. This model also suggested that TYK2 is predominantly necessary for IFNλ signaling in neutrophils but not epithelial cells (29). Additionally, JAK2 has been proposed as a transducer of IFNλ signaling (29, 33, 34). This is a feature shared with the IFN_γ_ receptor which engages both JAK1 and JAK2 to activate STAT1 homodimers to induce antiviral gene expression (35, 36). Whether ERK5 kinase activity selectively modulates IFN receptor density, distinct Janus kinase activity, or directly impacts STAT1 phosphorylation to regulate IFN-mediated gene expression warrants future study.

The loss of ULK1 has been shown to decrease the antiviral protection conferred by IFN_γ_ through decreased induction of gene expression (18). ULK1 deletion correlated with impaired ERK5 activation following IFN_γ_ stimulation. Thus, ERK5 could likely be a regulator of IFN_γ_-mediated gene expression. Treatment of murine embryonic fibroblasts and human fibrosarcoma cell lines with an ERK5 kinase inhibitor known to have off-target selectivity to BRD4 (XMD8-92), attenuated antiviral protection conferred by IFN_γ_ stimulation (18). BRD4 has been shown to mediate the transcription of ISGs, potentially confounding the specific requirement of ERK5 signaling in sustaining IFN_γ_ responses (37, 38). To overcome this, we utilized multiple ERK5 inhibitors with distinct chemical structures and distinct off-target profiles and also generated ERK5 null cells. Our study mechanistically demonstrates the specific requirement for ERK5 expression in the regulation of STAT1 activation in response to type III IFN. Whether ULK1 and ERK5 similarly control STAT1 activation or the activation of other transcription factors necessary to drive IFN_γ_-mediated gene expression remains to be determined.

The proinflammatory roles of ERK5 have been studied in the context of pathologies such as bacterial disease, ischemic injury, and cancer (16, 39). In these settings, ablation of ERK5 has proven to be beneficial. However, our study demonstrates that ERK5 constibutes to the protection against virus infections by enhancing type III IFN responses in epithelial cells. Together, these obseration demonstrate the need for a deeper understanding of the context in which ERK5 is activated and how it contributes to inflammatory control of distinct pathogens. A greater understanding of ERK5 activities will facilitate the creation of precise pharmacological interventions that will mitigate deleterious inflammation and uncontrolled cellular proliferation while preserving beneficial host protective responses.

## MATERIALS AND METHODS

### Cell lines, recombinant cytokines, and chemical treatments

Murine small intestinal epithelial cells (muINTEPI; InScreenex) and derived cell lines were cultured as previously described (6). Human A549 cells, HEK293FT cell lines were cultured in Dulbecco’s modified Eagle’s medium (DMEM) supplemented with 10% FBS, 2mM Glutamine, 100 U/ml Penicillin and 100 mg/ml Streptomycin and maintained at 37°C in 5% CO_2_. Non-targeted (H1) and *Mapk7*-deficient (*Mapk7* KO) cells were generated by CRISPR-Cas9 genome editing using previously described strategies (40). BHK21 and MDCK cells were obtained from BEI resources and grown in EMEM supplemented with 10% FBS, 2mM Glutamine, 100 U/ml Penicillin and 100 mg/ml Streptomycin and maintained at 37°C in 5% CO_2_. Cells were stimulated with human recombinant IFNλ3 (R&D Systems) or murine recombinant IFNβ (R&D Systems) and IFNλ3 (R&D Systems) at the indicated concentrations. ERK5 kinase inhibitors, XMD8-85, JWG-071, and AX15836 (Cayman Chemicals), were reconstituted in DMSO and used at the indicated concentrations.

### Small intestinal crypt and villi isolations

Murine small intestines from adult C57BL/6J male mice were removed and placed into ice cold PBS. Intestines were flushed twice with ice cold PBS and opened longitudinally. The mucus layer was gently removed by hand, and the small intestine was cut in half and designated as proximal and distal portions. Each portion was placed in separate conical tubes with 50ml of ice cold 10mM PBS-EDTA and incubated on ice for 45 min-1 hr. To separate the villi and crypts from the underlying connective tissue, the tissue was placed into a 100cm dish with ice cold 10mM PBS-EDTA and gently scraped using a microscope slide. The solution containing crypts and villi was vigorously mixed by pipetting and filtered through a 70-μm mesh to obtain purified crypt fractions. Villi that remained were returned to PBS-EDTA solution and this procedure was repeated two to three times to enrich for crypt fractions. Remaining villi were then resuspended in fresh 10mM PBS-EDTA in a 50-ml conical tube. All fractions were spun down and the supernatant poured off. Aliquots of crypt and villi fractions were resuspended in fresh RIPA buffer supplemented with Complete Protease Inhibitor Cocktail (Roche) and PhosSTOP phosphatase inhibitors (Roche) for further protein extraction and western blotting. Murine husbandry and tissue harvest was done following approved Institutional Animal Care and Use Committee and Institutional Biosafety Committee protocols at The Ohio State University.

### Plasmids and Oligonucleotides

CRISPR-Cas9 plasmids; pRRL-H1-PURO (non-targeting) and pRRL-*Mapk7*-PURO was generated by cloning single-guide RNA (sgRNA) targeting *Mapk7* (5’-AAAGGACGAAACACCGCGACGTGACCTTTGACGTGGGTTTTAGAGCTAGAAATAGCAAG -3’) and *Mapk7* (5’-CTTGCTATTTCTAGCTCTAAAACCCACGTCAAAGGTCACGTCGCGGTGTTTCGTCCTTT 3’) into empty pRRL-Cas9-PURO plasmids as previously described (6, 41). Lentiviral particles were generated in 293FT cells for transduction of muINTEPI cells followed by stable selection of transduced cells with puromycin.

### Viral infections and quantification

Vesicular Stomatitis Virus expressing Green Fluorescent Protein (VSV-GFP) was a gift from Dr. Dominique Garcin (Université of Genève) and was expanded and quantified in BHK21 cells (42). VSV-GFP infection was done at the indicated multiplicity of infection (MOI) and infectivity was assessed qualitatively by epifluorescent microscopy (EVOS) and crystal violet uptake assay, or quantitatively by flow cytometric analysis using a BD FACS CANTO II analyzer. VSV-GFP infectious virus production was measured by plaque assay in BHK21 cells. Influenza A virus (A/Puerto Rico/8/1934 (H1N1) tagged with green fluorescent protein (IAV-GFP) was grown in embryonated chicken eggs as previously described (43). IAV-GFP viral titers (TCID50) were determined in MDCK cells. A549 infections with IAV were conducted at an MOI 5 based on TCID50. Cell infectivity was assessed by flow cytometric analysis using a BD FACS CANTO II analyzer.

### Crystal violet uptake assay

Cells were pre-treated with ERK5 kinase inhibitors and/or recombinant IFN as indicated in figure legends. Cells were then infected with VSV-GFP at a multiplicity of infection (MOI) of 3. Following 24 h of infection, the culture medium was removed, cells were fixed with 4% PFA in PBS for 30 minutes at room temperature. Cells were washed with PBS and stained with Crystal violet stain (0.4% w/v) in 20% methanol for 30 minutes at room temperature. Plates were rinsed with distilled water to remove excess crystal violet stained and dried. Plates images were acquired using a ChemiDoc Touch (BioRad) imaging system. Distaining of cells was done by incubation with 1 ml of methanol for 30 minutes at room temperature with gentle shaking. A 50 µL aliquot of the distaining solution was transferred to clear bottom, low binding 96 well plate, and absorbance was measured at 595 nm.

### RNA extraction and quantification of gene expression

Total RNA was extracted from cells at the indicated time points post stimulation and/or infection using the NucleoSpin RNA extraction kit (Macherey-Nagel) as directed by manufacturer. cDNA synthesis was performed using iScript cDNA synthesis kit (BioRad) according to the manufacturer guidelines. Relative quantification of mRNA was done by quantitative PCR (qPCR) using a BioRad CFX-384 thermal cycler using iTAQ reagents (BioRad). Primers and probes used for qPCR assays are indicated in Supplementary Table 1.

### RNA sequencing, data processing, and analysis

Total RNA was extracted from muINTEPI cells seeded in 12-well plates and pre-treated with 10 μM of ERK5 kinase inhibitor, XDM8-85, for two hours prior to stimulation with 50 ng/ml of murine recombinant IFNλ3 for 24 hrs. Nucleic acid quantification, poly-A enriched library preparation, QC, and sequencing was carried out by Azenta. Samples had an average read depth of 40 million. Raw RNA-seq data quality control, alignment to mm10 genome (STAR), and count generation (HTSeq) was done using the Partek Flow bioinformatics software. Differential gene expression analysis was done using DESeq2 using Partek Flow. Expression heatmap and Venn diagram of DEG, and enrichment bubble plot were created with R (v2023.03.0+386) using libraries ‘ggplot2’ (v3.5.0) and ‘ggvenn’ (v0.1.10). Pathway enrichment analysis was performed with Enrichr (https://maayanlab.cloud/Enrichr/).

### Western blot analysis

Whole cell lysates were prepared with RIPA buffer (10 mM Tris-Cl (pH 8.0), 1 mM EDTA, 0.5 mM EGTA, 1% Triton X-100, 0.1% sodium deoxycholate, 0.1% SDS, 140 mM NaCl) supplemented with 1X protease and phosphatase inhibitor cocktail (Halt, Pierce) and Benzonase (Sigma). Protein concentrations were measured using the BCA Protein Assay Kit (Pierce) following manufacturer’s guideline. Proteins lysates were denatured with DTT and boiled prior to separation by SDS-PAGE and transferred onto PVDF membranes (Bio-Rad). Primary antibody incubations (Supplementary Table 2) were done overnight in 3% non-fat milk in Tris-buffered saline/Tween 20 followed by incubation with species-specific HRP-conjugated secondary antibodies (1:10,000; Jackson ImmunoResearch). Electrochemiluminescent image acquisition was done using a ChemiDoc Touch (BioRad).

### Protein sample preparation for Capillary-LC/MS/MS

Sample preparation and capillary-liquid chromatography-nanospray tandem mass spectrometry (Capillary-LC/MS/MS) conducted by the Mass Spectrometry & Proteomics (CCIC-MS&P) Shared Resource at The Ohio State University. muINTEPI cells pre-treated with 10 μM of ERK5 kinase inhibitor, XDM8-85, for two hours prior to stimulation with 50 ng/ml of murine recombinant IFNλ3 for an additional 24 hrs. Whole cell lysates were made in RIPA buffer (Cell Signaling Technologies). 20 µL of 10% SDS in 50mM Triethylamonium bicarbonate (TEAB) were added to 20 µL of protein samples (final concentration of SDS 5%). 5 µL of 50 mM ABC containing 5 µg/µL DTT were added prior to sample incubation at 65°C for 15 min followed by the addition of 5 µL of 50 mM ABC + 15 µg/µL iodoacetamide and incubation at RT for 15 min in the dark. Samples were acidified by adding 12% phosphoric acid (1:10 v/v acid to sample). 265uL of 1M TEAB /MeOH (10:90 v/v) was added and samples were loaded to S-trap for further washes. Samples were centrifuged at 4000 x g for 3 min (4°C) to remove supernatant. The trap was washed 3-6 times with 150uL of wash solution (1M TEAB /MeOH (10:90 v/v)). After the final wash, samples were digested with sequencing grade trypsin dissolved in 50mM TEAB with an O/N incubation at 37°C. Peptides were eluted with 40 µL of 50mM TEAB, 0.1% FA and 0.1% FA in Acetonitrile (50:50), sequentially. The sample was pooled together and dried in a vacufuge and resuspended in 20 µL of 50 mM acetic acid. Peptide concentration was determined by nanodrop (A280nm).

### Liquid chromatography-nanospray tandem mass spectrometry

Capillary-LC/MS/MS of protein identification was performed using a Thermo Scientific orbitrap Fusion mass spectrometer equipped with a nanospray FAIMS Pro™ Sources operated in positive ion mode. Samples (6.4 µL) were separated on an easy spray nano column (PepmapTM RSLC, C18 3µ 100A, 75µm X150mm Thermo Scientific) using a 2D RSLC HPLC system (Thermo Scientific). Each sample was injected into the µ-Precolumn Cartridge (Thermo Scientific) and desalted with 0.1% Formic Acid in water for 5 min. The injector port was switched to inject, and the peptides were eluted off the trap onto the column. Mobile phase A was 0.1% Formic Acid in water and acetonitrile (with 0.1% formic acid) was used as mobile phase B. Flow rate was set at 300 nL/min. mobile phase B was increased from 2% to 16% in 105 min and then increased from 16-25% in 10 min and again from 25-85% in 1 min and then kept at 95% for another 4 min before being brought back quickly to 2% in 1 min. The column was equilibrated at 2% of mobile phase B (or 98% A) for 15 min before the next sample injection.

MS/MS data was acquired with a spray voltage of 1.95 KV and a capillary temperature of 305 °C. The scan sequence of the mass spectrometer was based on the preview mode data dependent TopSpeed™ method: the analysis was programmed for a full scan recorded between m/z 375-1500 and a MS/MS scan to generate product ion spectra to determine amino acid sequence in consecutive scans starting from the most abundant peaks in the spectrum in the next 3 seconds. To achieve high mass accuracy MS determination, the full scan was performed at FT mode and the resolution was set at 120,000 with internal mass calibration. Three compensation voltage (cv=-50, -65 and -80v) were used for samples acquisition. The AGC Target ion number for FT full scan was set at 4 x 105 ions, maximum ion injection time was set at 50 ms and micro scan number was set at 1. MSn was performed using HCD in ion trap mode to ensure the highest signal intensity of MSn spectra. The HCD collision energy was set at 32%. The AGC Target ion number for ion trap MSn scan was set at 3.0E4 ions, maximum ion injection time was set at 35 ms and micro scan number was set at 1. Dynamic exclusion is enabled with a repeat count of 1 within 60s and a low mass width and high mass width of 10ppm.

Raw data were searched using Mascot Daemon by Matrix Science version 2.7.0 (Boston, MA) via ProteomeDiscoverer (version 2.4 Thermo Scientific,) and the database searched against the most recent Uniprot databases. The mass accuracy of the precursor ions was set to 10ppm, accidental pick of 1 13C peaks was included in the search. Fragment mass tolerance was set to 0.5 Da. Carbamidomethylation (Cys) is used as a fixed modification and considered variable modifications were oxidation (Met) and deamidation (N and Q). Four missed cleavages for the enzyme were permitted. The false discovery rate (FDR) by searching a decoy database and peptides were filtered at 1% FDR. Proteins identified with at least two unique peptides were considered as reliable identification. Any modified peptides are manually checked for validation.

## Acknowledgements

This work was supported in part by the National Institutes of Health under grant numbers R35 GM150806 (A.F.) and P30 CA016058 (The Ohio State University Comprehensive Cancer Center), The Ohio State University Award for Advancing Research in Infection and Immunity (A.F.), and the Ohio State University Center for Clinical and Translational Science voucher program under grant number RUL1TR02733. M.M.M is supported in part by DP2 CA271361 and Pew Biomedical Scholar Award. A.K is supported in part by Pelotonia Graduate Fellowship award.

The content is solely the responsibility of the authors and does not necessarily represent the views of the funding agencies.

## Author contributions

Investigation and formal analysis, H.B., D.S.N, M.I.M., A.K., A.B., A.K. and A.F.; conceptualization, D.S.N., A.K., and A.F.; methodology, H.B., P.D., and P.C.; data curation, H.B., P.C., A.F.; writing, A.F.; supervision, M.M.M, J.S.Y., and A.F.; funding acquisition, A.F.

## FIGURE LEGENDS

**Figure S1. Supplementary data related to Figure 1.**
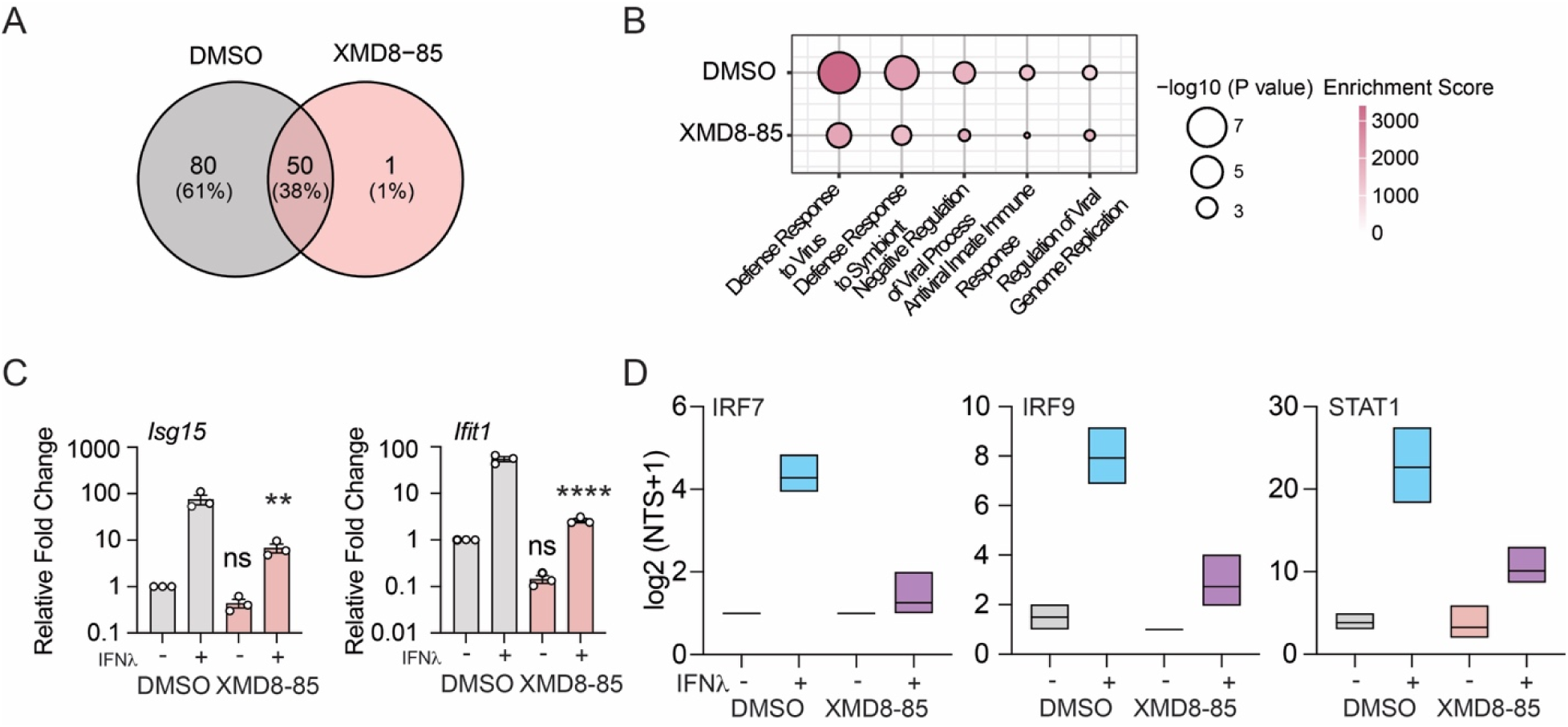
(A) Venn diagram of overlap of 131 DEG identified across DMSO or XMD8-85 IFN-treated cells relative to mock-stimulated cells. (B) Gene ontology enrichment of 131 DEG following IFN stimulation. Bubble size indicates significance (-log10 p-value) and color indicates enrichments score. (C) Relative expression of *Ifit1* and *Isg15* mRNA. Relative expression was normalized to vehicle treatment and *Chmp2a* expression (value 1). Data is representative of the average of three individual replicates and error bars represent ±SEM. (D) Protein expression of IFN-inducible transcription factors associated with ISG transactivation. Box and whisker plots represent the spectral counts corresponding to IRF7, IRF9, and STAT1 detected by LC-MS/MS in cells pretreated with vehicle or ERK5 inhibitor followed by IFNλ3 stimulation.

**Figure S2. Supplementary data related to Figure 3.**
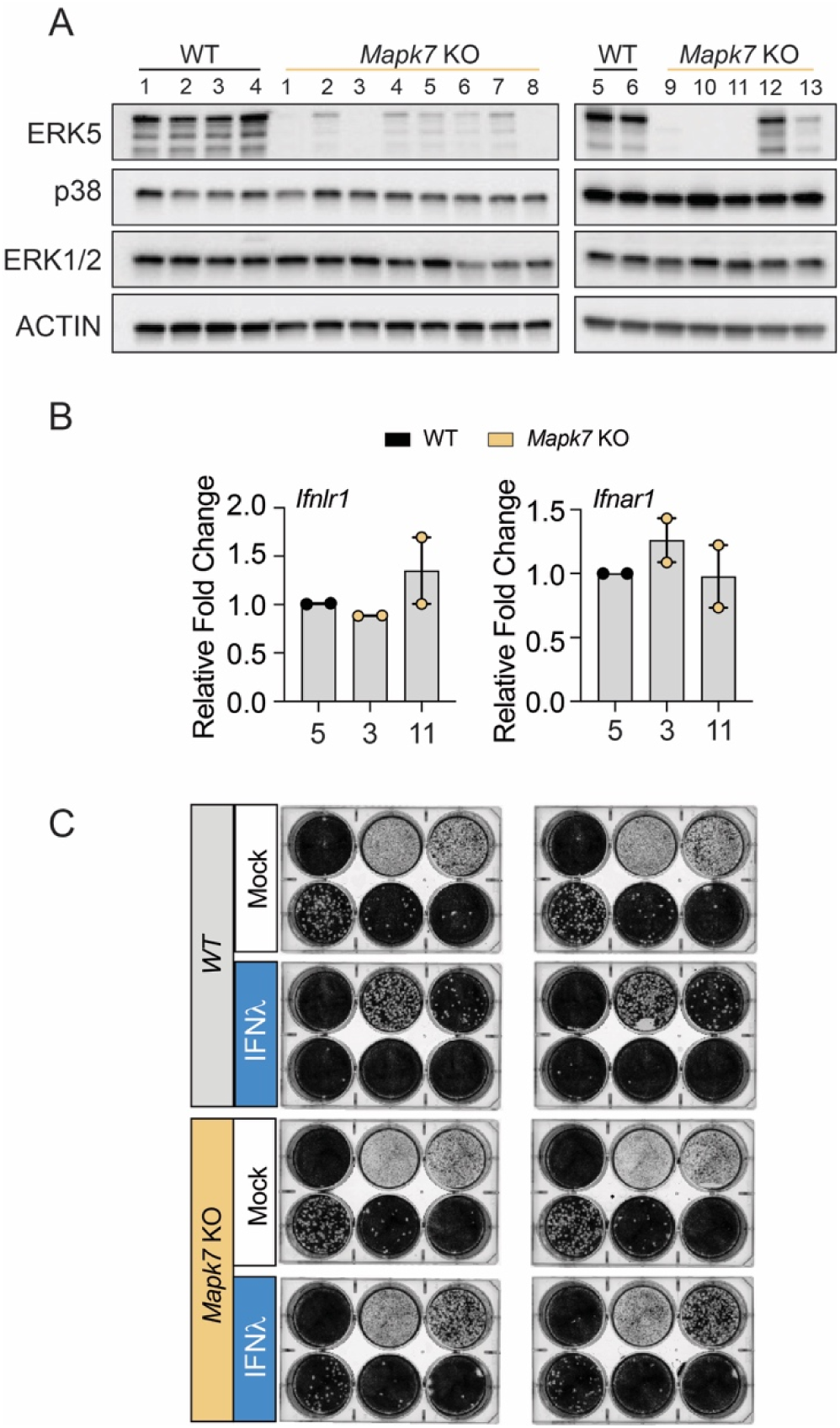
(A) Validation of ERK5 deletion across muINTEPI Clones. Total protein lysates from clonal cells transduced with Cas9 and non-targeting sgRNA (WT) or Mapk7-targeting sgRNA were probed for expression of ERK5, ERK1/2, p38, and Actin. (B) Expression of IFN receptors across *Erk5* targeted cells. Relative expression of *Ifnlr1* (left) and *Ifnar1* (right) mRNA. Relative expression was normalized to wild-type (H1) and *Hprt1* expression (value 1). Data is representative of the average of two individual replicates and error bars represent ±SEM. (C) Representative plaque assay quantification from infectious virus production in WT and *Mapk7* KO cell infected with VSV-GFP. Cells were treated IFNλ (100 ng/ml) for 24 hrs prior to preparation of whole cell lysates. Data representative of 4 independent experiments.

