## Supplementary Table 1 for "ERK5 Signaling is Required for Type III IFN-mediated Mucosal Antiviral Responses"

### TABLE EV1. Reagent Table

| **REAGENT** | **VENDOR/SOURCE** | **IDENTIFIER** |
| --- | --- | --- |
| **Antibodies** | | |
| Donkey Anti-Mouse IgG (HRP-Conjugated) | Jackson ImmunoResearch | RRID: AB_2340770 |
| Donkey Anti-Rabbit IgG (HRP-Conjugated) | Jackson ImmunoResearch | RRID: AB_10015282 |
| ISG15 (F-9) | Santa Cruz | RRID: AB_2126308 |
| Actin-HRP | Santa Cruz | RRID: AB_626632 |
| STAT1, phospho (Tyr701) | Cell Signaling Technology | RRID: AB_561284 |
| STAT1 (D1K9Y) | Cell Signaling Technology | RRID: AB_2737027 |
| ERK5 | Cell Signaling Technology | RRID: AB_330491 |
| p44/42 MAPK (Erk1/2) | Cell Signaling Technology | RRID:AB_390779 |
| SAPK/JNK | Cell Signaling Technology | RRID:AB_2250373 |
| p38 MAPK (D13E1) XP | Cell Signaling Technology | RRID:AB_10999090 |
| Phospho-ERK5 (Thr218/Tyr220) | Cell Signaling Technology | RRID:AB_2140424 |
| IFITM1 | Cell Signaling Technology | RRID:AB_2798126 |
| IFITM3 | ProteinTech | RRID:AB_2295684 |
| RIG-I (Alme-1) | AdipoGen Life Sciences | RRID:AB_2490189 |
| IFIT3 (B-7) | Santa Cruz | RRID:AB_2857847 |
| **Bacterial and Virus Strains** | | |
| VSV-GFP | Dominique Garcin | Hausmann et al., 2008 |
| Influenza A-GFP Strain: A/Puerto Rico/8/1934 (H1N1) | Adolfo Garcia-Sastre | Manicassamy et al., 2010 |
| Stellar Competent Cells *E. coli* HST08 strain | Takara | Cat# 636763 |
| **Chemicals, Peptides, and Recombinant Proteins** | | |
| Human recombinant IFNλ3 | R&D Systems | 5259-IL-025 |
| Murine recombinant IFNβ | R&D Systems | 8234-MB-010/CF |
| Murine recombinant IFNλ3 | R&D Systems | 1789-ML-025/CF |
| Mirus Bio TransIT-X2 | Mirus Bio LLC | MIR6000 |
| XMD8-85 | Cayman Chemicals | 29662  Batch: 01581666-2 |
| JWG-071 | Sigma-Adrich | SML2541 |
| AX15836 | Tocris | Cat. No. 5843/10 |
| 100x Halt protease and phosphatase inhibitor | Thermo Fisher Scientific | 78420 |
| PhosSTOP, phosphatase inhibitor | Sigma-Aldrich | 4906845001 |
| cOmplete Protease Inhibitor Cocktail | Sigma-Aldrich | 046931322001 |
| Benzonase®, Nuclease | Sigma-Aldrich | E1014-25KU |
| **Commercial Assays** | | |
| NucleoSpin RNA II | Macherey-Nagel | 740955.250 |
| iSCRIPT cDNA Synthesis Kit | Bio-Rad | 1708891 |
| SsoAdvanced Universal Probes Supermix | Bio-Rad | 1725281 |
| TaqMan Universal Master Mix II, no UNG | Thermo Fisher Scientific | 4440048 |
| Pierce BCA Protein Assay | Thermo Fisher Scientific | PI23227 |
| **Cell Lines** | | |
| muINTEPI | InScreenex | Cat# INS-CI-1007 |
| A549 | ATCC | Cat# CRL-1593.2, RRID: CVCL_0007 |
| *H1* muINTEPI | This Paper | N/A |
| *Mapk7* KO muINTEPI cells | This Paper | N/A |
| 293FT | ATCC | CRL-1573 |
| MDCK | BEI | NR-2628 |
| BHK21 | ATCC | CCL-10 |
| PH5CH8 | Michael Gale Jr. | Forero et al., 2019 |
| **Oligonucleotides** | | |
| *Mapk7* sgRNA S: 5’-AAAGGACGAAACACCGCGACGTGACCTTTGACGTGGGTTTTAGAGCTAGAAATAGCAAG -3’ | Integrated DNA technologies | N/A |
| *Mapk7* sgRNA AS: 5’- CTTGCTATTTCTAGCTCTAAAACCCACGTCAAAGGTCACGTCGCGGTGTTTCGTCCTTT-3’ | Integrated DNA technologies | N/A |
| *Chmp2a* F: 5’- AGACGCCAGAGGAACTACTTC-3' R: 5’- ACCAGGTCTTTTGCCATGATTC-3' | Integrated DNA technologies | N/A |
| *Ifi204* F: 5’- CCAGTCACCAATACTCCACAGC-3' R: 5’- CTCTGAGTGGAGAACAGCACCT-3' | Integrated DNA technologies | N/A |
| *Ifi4*: F: 5'- ATGCACTCTTCTGAGCTGGTGG-3' R: 5’- TCAGATCCAGGCTATCCACGTG-3' | Integrated DNA technologies | N/A |
| *Ifit1* F: 5’- CTGAGATGTCACTTCACATGGAA-3' R: GTGCATCCCCAATGGGTTCT-3' | Integrated DNA technologies | N/A |
| *Isg15* F: 5’- GGTGTCCGTGACTAACTCCAT-3' R: 5’- TGGAAAGGGTAAGACCGTCCT-3' | Integrated DNA technologies | N/A |
| *Oas2* F: 5’- CACCAAAGTCCTGAAGACCGTC-3' R: 5'- AGAGTCGTAACTCTCCAGCGAG-3' | Integrated DNA technologies | N/A |
| *IFITM1* F: 5'-GCACAACCAGGAACATGAG-3' R: 5'-CAGCACCAGTTCAAGAAGAG-3' | Integrated DNA technologies | N/A |
| *IFITM3* F: 5'-ATCACACTGTCCAAACCTTCT-3' R: 5'-GTGGATCACGGTGGACGTC-3' | Integrated DNA technologies | N/A |
| *ISG15* F: 5’- CGCAGATCACCCAGAAGAT-3'  R: 5’-CAGAGGTTCGTCGCATTTG-3' | Integrated DNA technologies | N/A |
| *HPRT1* F: 5’- CATTATGCTGAGGATTTGGAAAGG-3'  R: 5’- CTTGAGCACACAGAGGGCTACA-3' | Integrated DNA technologies | N/A |
| *Il28ra* Probe: 5’- /56-FAM/TGTGTCCCC/ZEN/TGATGTTGCCTGAAG/3IABkFQ/-3’ | Integrated DNA technologies | Mm.PT.58.10781457 |
| *Ifnar1* Probe: 5’- /56-FAM/AGCCTCCCC/ZEN/GCAGTCTTGATGAG/3IABkFQ/-3’ | Integrated DNA technologies | Mm.PT.58.33360098 |
| *Hprt* Probe: 5’- /56-FAM/CTTGCTGGT/ZEN/GAAAAGGACCTCTCGAA/3IABkFQ/-3’ | Integrated DNA technologies | Mm.PT.39a.22214828 |
| **Plasmids** |  |  |
| pRRL-H1-PURO | Daniel Stetson | Froggatt et al., 2021 |
| pRRL-Mapk7-PURO | This paper | Gray et al., 2015 |
| **Software** |  |  |
| GraphPad Prism 9 | GraphPad Software | RRID: SCR_002798 |
| R Studio v2023.03.0+386 | Rstudio | RRID: SCR_000432 |
| ggplot2 v3.5.0 | Wickham, 2016 | https://github.com/tidyverse/ggplot2 |
| ggvenn v0.1.10 | Yan Linlin | https://github.com/yanlinlin82/ggvenn |
| EnrichR | Chen et al., 2013 | RRID: SCR_001575;  https://maayanlab.cloud/Enrichr/ |
| Partek Flow | Partek | https://www.partek.com/partek-flow/ |
| CFX Maestro | BioRad | Catalog number: 12013758 |
